# Isolation and characterization of broad-range *Staphylococcus epidermidis* sepunaviruses

**DOI:** 10.1101/2025.10.14.682026

**Authors:** Carlos Valdivia, Callum O. Rimmer, Jonathan C. Thomas, David Negus, Pilar Domingo-Calap

## Abstract

*Staphylococcus epidermidis* is a major contributor to biofilm-associated infections, bacteraemia, and sepsis in humans. Additionally, it is an important veterinary pathogen. The spread of multi-drug resistant *S. epidermidis* (MRSE) poses an even greater challenge, requiring the development of antibiotic-alternative approaches. Here, we isolated four new broad-range phages capable of infecting a large panel of 65 clinical and veterinary isolates of *S. epidermidis*. Phages with the broadest host range produced plaques on 37% of the hosts tested and halos on an additional 40%. These phages belonged to the *Sepunavirus* genus, as supported by their morphology and genome analysis. A genome-wide association study identified a significant correlation between resistance to phage lysis and the presence of the *sbcCD* complex and the *ica* operon, highlighting the protective role of biofilms against the phages isolated in this study. Furthermore, distinct phage-encoded methyltransferases identified in the phage genomes may contribute to differences in host ranges. This study advances our knowledge on the diversity and characteristics of *S. epidermidis* phages, which could be considered as interesting tools for phage therapeutics.

**Repositories:** Phage genomes are available under Genbank accession numbers PV426907-PV426910. Bacterial genome sequences determined as part of this study, and all Bakta annotations, are available via Figshare (doi:10.6084/m9.figshare.29066417 and doi:10.6084/m9.figshare.29079968 respectively).

## Introduction

The emergence and rapid spread of multi-drug resistant (MDR) bacteria worldwide has compromised the effectiveness of antibiotics. This phenomenon has led to the emergence of bacterial isolates resistant to all available antibiotics, complicating treatment and leaving healthcare providers with few viable options^1,2^. In 2019, 5 million deaths were associated with MDR bacteria, with 1.27 million directly attributable to them^3^. Without alternative treatments, infections caused by MDR bacteria could result in up to 10 million deaths annually by 2050^4^. Aside from healthcare, other fields such as agriculture, livestock farming, and various industries also struggle with bacterial infections and contaminations, posing challenges that impact humans, animals, and ecosystems alike^5^.

Bacteria develop resistance through various mechanisms, including modification of the antibiotic target, degradation of the antibiotic, or blocking access to its target. One of the most significant contributors to antibiotic resistance is the formation of biofilms^6^. Biofilms are structures in which bacteria aggregate, embedding themselves in a self-produced polysaccharide matrix rich in proteins and extracellular DNA. The use of indwelling medical devices, such as feeding catheters, can facilitate biofilm-associated infections. Bacteria from a contaminated device can migrate into the bloodstream, leading to acute late-onset sepsis^7^. Some biofilm-forming bacteria like *Staphylococcus epidermidis* combine this trait with a widespread colonization of human epithelia^8,9^. This makes *S. epidermidis* a significant nosocomial pathogen and one of the most frequent causes of bacteraemia and sepsis, particularly in neonates and immunocompromised patients^7,10,11^. Neonates are rapidly colonized by *S. epidermidis*, which helps prevent colonization by more virulent bacteria like *S. aureus*^12^. However, neonates in intensive care units (ICUs) frequently acquire MDR strains of *S. epidermidis,* resulting in bacteraemia. Although mortality rates are relatively low, these infections lead to prolonged hospital stays and elevated morbidity, also being associated with post-inflammatory sequelae^11,13^.

This situation is further complicated by the spread of MDR isolates, including methicillin-resistant *Staphylococcus epidermidis* (MRSE)^14–16^. Initially, methicillin was the first-line treatment for *S. epidermidis* infections; however, resistance has now been documented in up to 75–90% of hospital isolates^11^. Biofilms reduce the efficacy of various antimicrobial agents, including vancomycin, which is now the standard therapy for MRSE^17^. *S. epidermidis* isolates commonly accumulate resistance to other antibiotics like rifampicin, fluoroquinolones, gentamicin, tetracycline, chloramphenicol, erythromycin, clindamycin, sulfonamides, and streptogramins. The high costs and lengthy development process for new antimicrobials, combined with the rapid emergence of resistance that shortens their effective lifespan, significantly reduce the economic incentives for pharmaceutical companies^18–20^.

*S. epidermidis* also poses significant threats in veterinary medicine. Most staphylococcal infections in animals can be managed with antibiotics selected based on antibiogram results^21^. However, pets and livestock are frequently colonized by MRSE strains that also show resistance to multiple antibiotics^21^. In ruminants, *S. epidermidis* is the predominant etiological agent of intramammary infections and is also implicated in various diseases in companion animals^22,23^. Mastitis is particularly problematic, as infections frequently remain subclinical yet persistent, leading to tissue damage and reduced milk quality^24,25^. Also, the zoonotic potential is noteworthy, as bacteria and their toxins can be transferred through milk, highlighting the need for antibiotic-alternative methods to manage these infections^24^. The success rate of antibiotic treatment for mastitis is highly variable, with non-curative outcomes often reported^26^. Overuse of antibiotics in bovine mastitis management exacerbates the development of antimicrobial resistance in *S. epidermidis* and other bacterial species, and also raises public concerns about antibiotic residues in the food supply^25^.

To address these challenges there is a need for therapeutic alternatives to combat MRSE infections. Promising approaches include probiotics, antibodies, antimicrobial peptides, and bacteriocins^27,28^. In addition, phage therapy has emerged as one of the most promising options. Bacteriophages, or phages, are viruses that infect bacteria. They represent the most numerous biological entities on Earth, showing great diversity regarding traits like morphology, specificity or genomic traits^29,30^. Due to their ability to infect and often kill their bacterial hosts, their use as antibacterial agents was proposed soon after their discovery^31^. However, their application needs to be carefully studied, addressing matters such as host specificity, lifestyle, and the potential to mobilise host derived genes associated with antibiotic resistance and virulence^32,33^. Bacteria have evolved a variety of defence systems to protect themselves from phage infection. The most common are restriction-modification (RM) systems. Here, differences in the epigenetic pattern, generally methylation, allow external genetic material to be considered foreign. Newly created bacterial DNA is constantly modified by proteins such as methyltransferases (MTases), generating a fingerprint. When a genetic fragment is considered foreign, bacterial restriction enzymes cleave them to protect bacteria from potentially harmful genetic elements^34,35^. Other defence systems can be adaptive or sequence-dependent, such as CRISPR-Cas or others^36^. In turn, phages have evolved a myriad of strategies to overcome these defences, like introducing mutations in target sequences, mimicking methylation patterns, or inhibiting the proteins involved in the defence, like the anti-CRISPR (Acr) proteins^37^. This constant struggle between bacteria and their phages results in an arms race, allowing phages to evade defence systems and modulating bacterial populations^36^. The existing diversity in defence and anti-defence systems makes it necessary to understand the correlation between host defence systems and susceptibility to specific phages to select viruses for phage therapy.

Here, we isolated and characterized four new broad-range *S. epidermidis* phages, showing strong lytic activity against a large collection of clinical and veterinary *S. epidermidis* isolates. Through bioinformatic analysis of the bacterial and phage genomes, we identified genes and defence systems potentially linked to phage resistance, as well as viral genes that may help phages overcome bacterial defences.

## Materials and methods

### Bacterial isolates

*S. epidermidis* ATCC 12228 and ATCC 35984 were obtained from the Spanish Type Culture Collection (CECT). In addition, a collection of 65 *S epidermidis* isolated from a range of sources were tested for susceptibility to phage lysis (Table S1). This included reference strains, isolates from veterinary samples and clinical samples. A reference *S. aureus* strain, ATCC 6538, was also included. Bacterial overnight cultures were grown from glycerol stocks stored at -70 °C in tryptone soy broth (TSB) at 37 °C with gentle agitation.

### Phages

Phages were isolated using the reference *S. epidermidis* ATCC 12228 as a primary host. For phage hunting, samples from waste-water treatment plants in the area of Valencia (Spain) were tested as described previously^38^. Individual plaques were picked and serially transferred three times to ensure purity. Phages were then propagated following overnight incubation at 37°C and 250 rpm to obtain high-titer lysates (>10^8^ PFU/mL), which were then stored at -70 °C.

### Transmission electron microscopy

Carbon-coated copper grids were glow-discharged (30 sec, 7.2V, Bal-Tec MED 020 Coating System) and placed on high-titer sample drops for 10 min. After two distilled water washes, grids were stained with 2% uranyl acetate for 5 min, excess fluid was removed, and grids were air-dried. Imaging was conducted with an FEI Tecnai G2 Spirit transmission electron microscope at 80 kV (ThermoFisher, Oregon, USA).

### Phage sequencing, genomic characterization and phylogenetic analysis

Phage lysates over 10^8^ PFU/mL were used for genome extraction after degradation of non-encapsidated DNA followed by capsid degradation as described previously^39^. Concentration and purification of phage DNA was achieved using the Maxwell® RSC (Promega) instrument. Sequencing libraries were prepared using the Nextera XT DNA kit and sequenced on an Illumina MiSeq instrument using 2×250 paired-end v3 chemistry. The quality of paired-end reads was assessed using FastQC v0.12.1. Acceptable reads were subsequently used as input for *de novo* assembly with Unicycler v0.5.0^40^. The assembly was then corrected with Pilon v1.24^41^ and overlapping ends were manually trimmed. Genomes are available under Genbank accession numbers PV426907-PV426910. Intergenomic similarity between the phages was assessed through VIRIDIC^42^. Viral family and lifestyle were predicted with PhaGCN^43^ and PhaTYP^44^. Full genome annotation was performed as recommended by using both Pharokka^45^ and Phold^46^. Where no gene product could be assigned, Foldseek^47^ was used. PhageRBPdetect v3^48^ was used to locate receptor binding proteins (RBPs) while proteins with potential depolymerase activity were specifically detected using PhageDPO^49^, DePP^50^ and Deposcope^51^. AntiCRISPR proteins (Acr) were analyzed with ACRFinder^52^ and AntiCRISPRdb v2.2.^37^ and the obtained sequences compared against the phage genomes with BLASTn^53^. Anti-defense systems were searched for with Defense Finder^54^. Palindromic sequences were detected using EMBOSS palindrome^55^ with default settings. To observe whole genome disposition and modularity, the R package gggenomes was used. For 3D modelling, potential oligo state was determined with SWISS-MODEL^56^, the 3D structure was modelled with AlphaFold 3^57^ and visualized with Chimera v.1.17.3^58^. Identity between 3D structures was assessed using a Pairwise comparison with DALI^59^.

Whole-genome phylogenetic analysis was performed using the genomes of the closest staphylophages according to BLASTn. After reordering and alignment using MAFFT v7.526.^60^, IQtree v2.3.5.^61^ was used to construct a maximum likelihood phylogeny with 1000 bootstrap replicates and default settings. The best tree was represented with ITOL v6^62^ after collapsing nodes with a bootstrap value below 90 and fixing the root in the midpoint.

### Bacterial sequencing and characterization

Details of the *S. epidermidis* isolates included in the host range assay can be found in table S1. Where whole genome sequencing data is available, isolates were sequenced using either Illumina or a hybrid Oxford Nanopore Technologies (ONT) MinION/Illumina approach, details in table S1. Coding sequences (CDS) and functional annotation were predicted using Bakta v1.9.3^63^ with default parameters. The annotated genomes were subsequently analyzed using Padloc v2.0.0^64^ and DefenseFinder v1.2.2^34^ to identify phage defence mechanisms, and CRISPRCasFinder v4.3.2 to retrieve CRISPR sequences throughout the genomes. All three programs were run using default parameters. Additionally, all sequenced genomes were examined with the online tool PHASTEST^65^ to locate prophage sequences. Sequences classified as prophages regardless of their score and classification (intact, questionable or incomplete) were annotated using Pharokka^45^. The presence of defence systems (DF) and prophages was correlated with phage infectivity through differential probability of resistance (dPR), as described in previous work^66^. Briefly, dPR = p(S|P) – p(S|A), where S indicates susceptibility to a given phage, P indicates the presence of a defence system, and A indicates its absence. dPR was calculated for every defence system as classified by PADLOC and DefenseFinder and negative dPR were considered to correlate with phage resistance. Only DFs present and absent in at least 9 genomes were considered.

### Phage host range evaluation

Host range analysis was performed as described previously^67^. Briefly, semi-solid tryptone soy agar (TSA; 5 ml; 0.6% agar) supplemented with CaCl_2_ and MgCl_2_ (both at a final concentration of 5 mM) was aliquoted into sterile test tubes held at 45 °C. Each tube was then inoculated with an overnight culture of the prospective host strain, and gently swirled to mix the contents before being poured onto a TSA plate. The plate was gently swirled to ensure even distribution of top agar. Once set, aliquots of phage lysate (titer > 10^5^ PFU/mL) were spotted onto the top agar. Plates were incubated overnight at 37 °C. Next day, plates were inspected for lysis, with results recorded according to a modification of Haines *et al*^68^, including complete lysis, turbid lysis, or no visible plaques.

### Phage adsorption assay

Phages were exposed at a low MOI (multiplicity of infection, 0.001) to different bacterial strains at 0.2 OD_600_. This included *Escherichia coli* C IJ1862, obtained from Prof. James J. Bull, as negative control. For the initial time point, an aliquot of the mixture was immediately centrifuged at 14,000 rpm for 5 minutes to pellet the bacterial cells along with any adsorbed phages. The supernatant was then collected and titrated to determine the concentration of free (unadsorbed) phages. Another aliquot of the mixture was incubated at 37 °C with shaking at 200 rpm for 35 minutes, a duration selected based on preliminary experiments indicating maximal reduction in free phage titer for both phages. The process was repeated as with the initial time point. To assess statistical significance (*p* < 0.05) a Mann-Whitney U test was performed.

### Genome-wide association study

The pan-genome of the available bacterial genomes (49 out of 65 bacterial isolates had been whole-genome sequenced) was determined using Panaroo v1.3.3^69^ with strict clean mode. GFF3 files produced by Bakta were converted to Prokka format using the convert_refseq_to_prokka_gff.py script prior to running Panaroo. Isolate phenotypes were assigned depending on their susceptibility to each of the phages, and were considered susceptible if clear or partial lysis/turbid plaques were observed. Scoary v1.6.16^70^ was used to determine genome-wide associations (GWAS) between presence/absence of genes and susceptibility to each phage. Genes were considered significant after correcting for multiple test comparisons using the Bonferroni method^71^.

## Results

### Isolation of broad-range *S. epidermidis* phages

Phage hunting was performed using sewage water from Valencia (Spain), using a reference *S. epidermidis* strain (ATCC 12228) as a primary host. Four single plaques which had similar morphology were isolated, purified and amplified. High titer lysates were used to assess the host range over 65 isolates of *S. epidermidis* from different sources, and a reference *S. aureus* strain. The infectivity matrix revealed two main patterns (Figure 1). Phages vb_sep_Steph1 (Steph1), vb_sep_Steph3 (Steph3) and vb_sep_Steph4 (Steph4) exhibited a broad host range, lysing 25 isolates efficiently, and producing hazy lysis over at least another 25. On the other hand, phage vb_sep_Steph2 (Steph2) infected 12 isolates, and showed some effect over another seven isolates. None of the phages was able to infect the reference *S. aureus* isolate.

**Figure 1.**
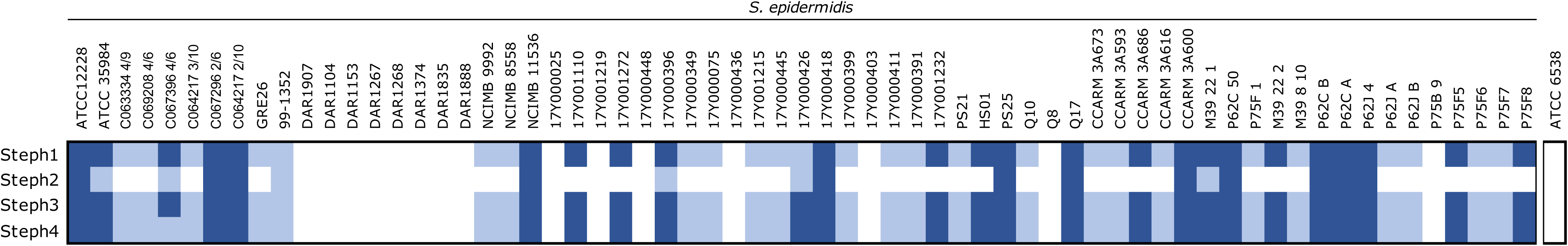
Infectivity matrix of the four *S. epidermidis* phages Steph1, Steph2, Steph3, and Steph4 over a panel of 66 *Staphylococcus* spp. isolates. Infectivity was determined by spot test, clear lysis (dark blue), turbid lysis (light blue), no lysis (white). *Staphylococcus aureus* ATCC 6538 is shown on the right.

### High similarity and modularity among the phage genomes

The four high-titer lysates were used for DNA extraction and sequenced. Bioinformatic analysis showed highly similar genomes (≈140 kbp, 28% GC content, minimum intergenomic similarity = 87.85%). All four phage genomes were predicted to have a virulent lifestyle and belonged to the family *Herelleviridae*, which was supported by electron microscopy analysis (Figure 2). The closest related phages were identified using BLASTn, with the most similar described phages being *Staphylococcus* phage BESEP4 (93.30% Intergenomic similarity to Steph4. Genbank <u>MT596501.1</u>) and *Staphylococcus* phage 110 (90.31% Intergenomic similarity to Steph3. Genbank <u>OQ448195.2</u>), both belonging to the *Sepunavirus* genus. Gene modularity showed a highly similar distribution of all genes (Figure 3). Alignment of the phage genomes and visual inspection identified a region of dissimilarity at approximately 42 kbp between phage Steph2 and the other three phages. At this location, phage Steph2 contained CDS0102, a predicted DNA methyltransferase (MTase) according to both sequence and structure-based annotators. No sequences similar to this protein were found in any of the other genomes. In the same location, Steph1, Steph3 and Steph4 shared a hypothetical protein absent in Steph2. Analysis of the 3D structure of both proteins (Figure 4) showed that, despite the lack of sequence similarity, the hypothetical protein in Steph1, Steph3 and Steph4 had a structure related to Steph2 MTase (up to 30% structural identity as homodimers), while sharing a 27 amino acid-long loop to which no structural equivalence could be found in Steph2. These CDSs were assigned the same classification as CDS0102 by Foldseek and were also detected as MTases by PHOLD. The loop was weakly associated to ribosomal proteins and detected in other staphylophages by HHPRED. Steph2 MTase was assigned site-specific (cytosine-N4-specific activity) activity based on Interpro search, and its classification as MTase was supported by PANTHER. In addition, both Steph2 and the other phages’ methyltransferases were detected as such by Interpro and HHPRED. Furthermore, the four DNA MTases were also detected as anti-defence systems by DefenseFinder. Another region at approximately 20 kb contained a shared hypothetical protein-encoding gene in Steph1, Steph3 and Steph4 whereas Steph2 had an unrelated gene encoding a hypothetical protein, but no role could be assigned to this CDS through sequence or structural associations.

**Figure 2.**
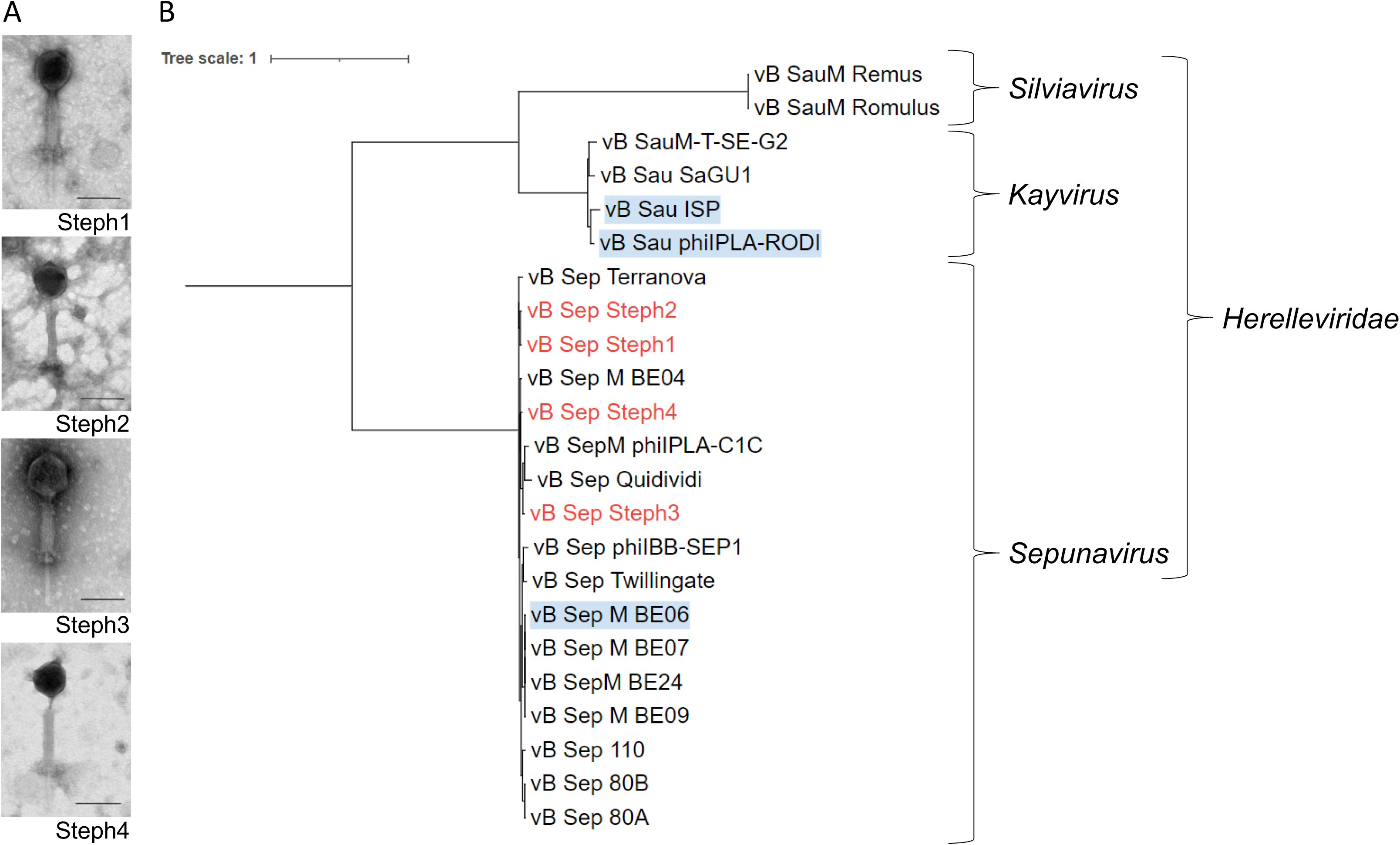
Taxonomic classification of *S. epidermidis* phages. A. Electron micrographs of phages showing myovirus morphology. Scale bar =100 nm. B. Phylogenetic tree of the four *S. epidermidis phages* Steph1, Steph2, Steph3, and Steph4 (marked in red) with closely related phage sequences belonging to the *Herelleviridae* family. The tree was constructed using maximum likelihood phylogeny with 1000 bootstrap replicates and rooted at the midpoint. Phages highlighted in blue indicate prior use in phage therapy.

**Figure 3.**
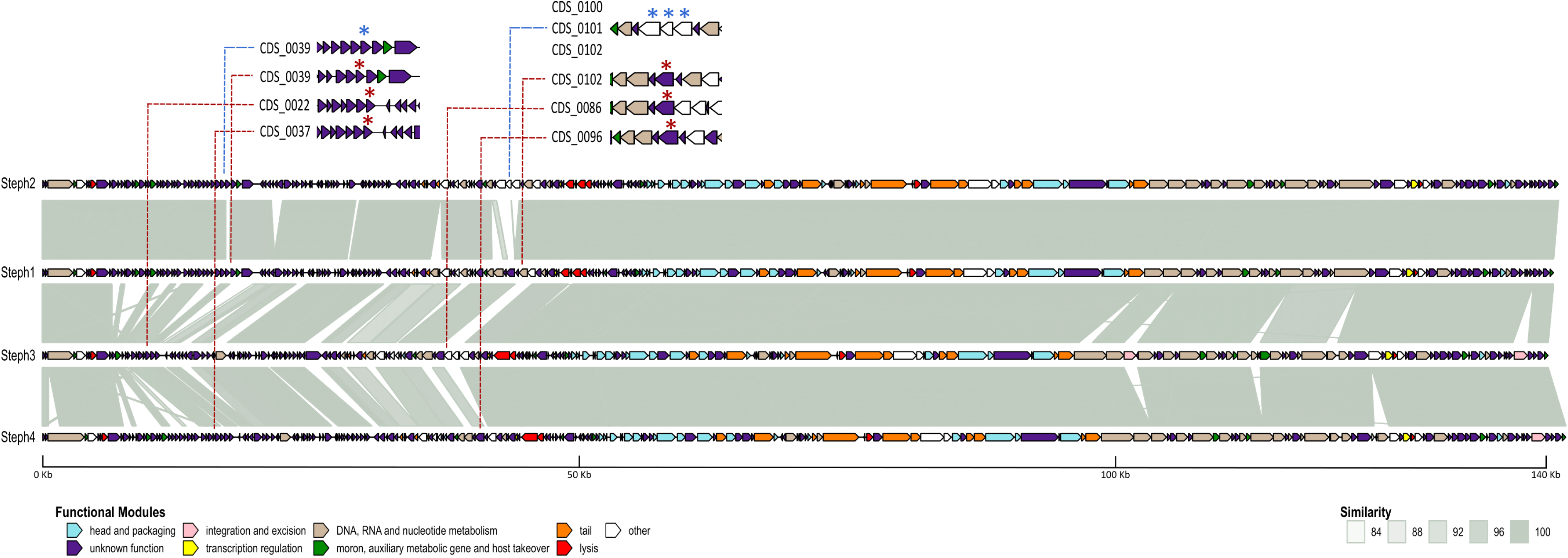
Comparative genomics of the *S. epidermidis* phages Steph1, Steph2, Steph3 and Steph4. Homology between genes was established based on BLASTn comparisons, similarity is shown in shades of grey. Functions of each annotated product are colored according to the legend.

**Figure 4.**
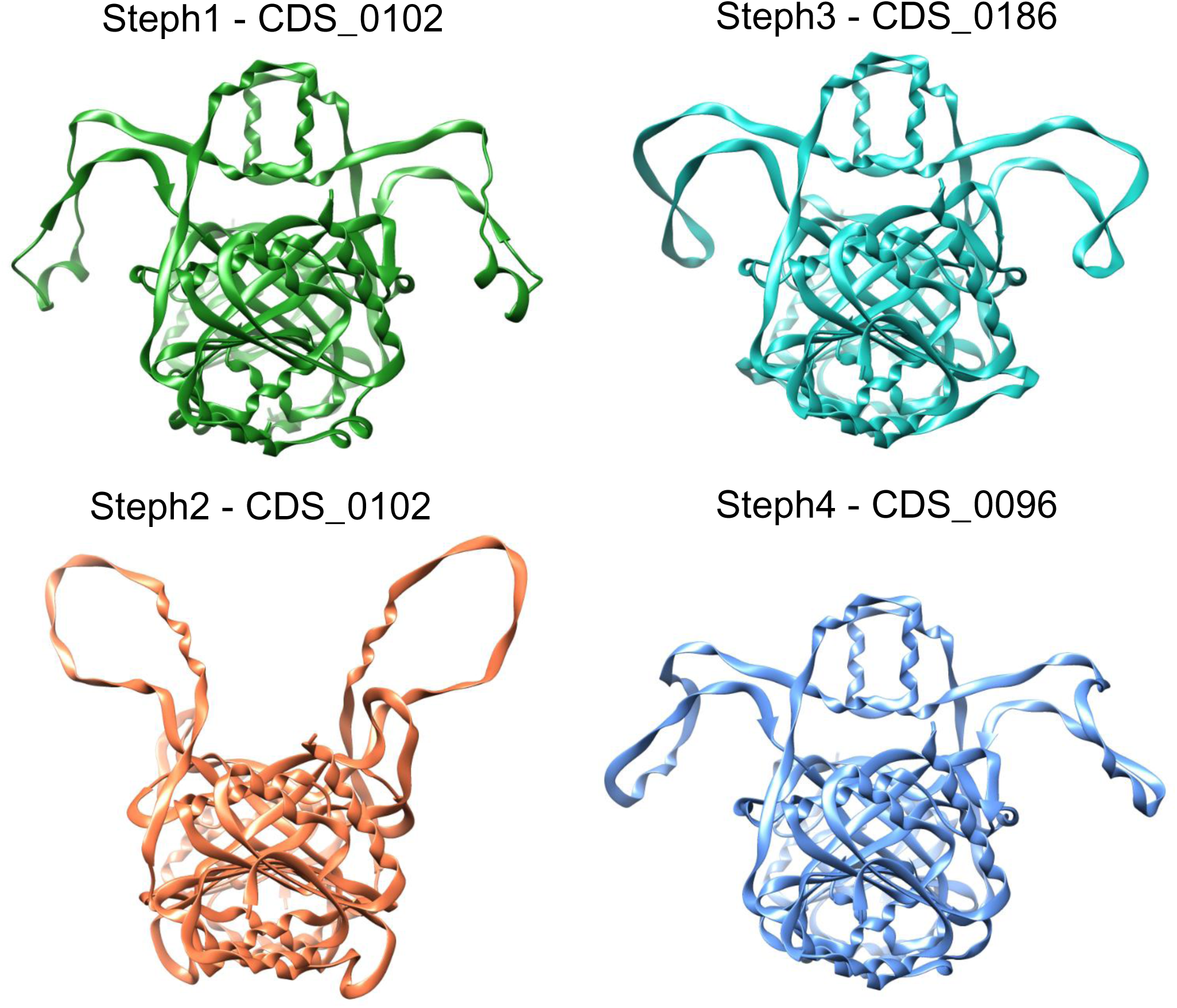
3D structures of the different MTases predicted using AlphaFold. The structures are presented as homodimers and visualized in Chimera, illustrating the similarities and differences in their 3D conformations.

Regarding proteins involved in recognition and adsorption, one single nucleotide polymorphism (SNP) located on CDS_0179, a virion structural protein, was found to differentiate Steph2 from the broad host range phages (Table S2). No further differences were observed between the broad-range phages and Steph2 with respect to recognition and adsorption proteins. All phages encoded 2 major tail proteins, 3 tail fiber proteins and 10 virion structural proteins aside from other elements of the tail, baseplate, and head. Also, each phage encoded two predicted RBPs, out of which one had putative depolymerase activity. Another CDS classified as a structural protein also had putative depolymerase activity.

### Determinants of bacterial traits on phage infectivity

The first approach to find determinants of phage infection was to evaluate the potential effect of the point mutation in CDS_0179 on the adsorption of phage Steph2. For this purpose, the adsorption of the phage was compared with that of the broad-spectrum representative phage Steph4. The tests were carried out on two strains of *S. epidermidis*, one susceptible to both phages and one in which Steph2 was unable to induce plaques. An *E. coli* strain was also included as a non-adsorptive control. The results obtained indicated that both phages adsorbed to the *S. epidermidis* strains, as opposed to the *E. coli* (Figure 5). Thus, the ability of Steph2 to bind without infecting suggested that post-entry mechanisms were involved in the observed difference in host range.

**Figure 5.**
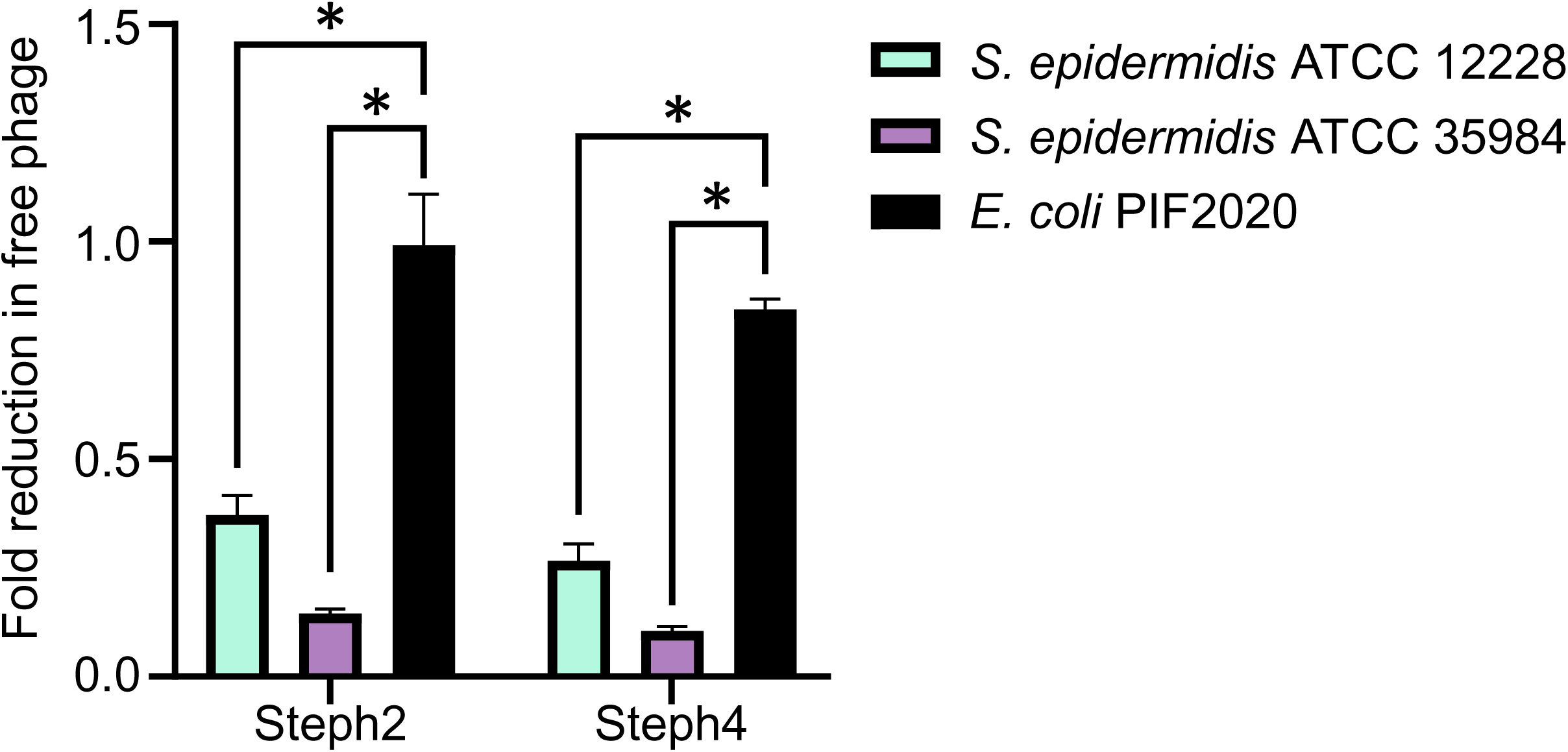
Fold reduction in free phage titer following adsorption. Data represents the mean of three independent experiments, each comprising three technical replicates. Error bars indicate the standard error of the mean. Statistical significance was assessed using the Mann-Whitney U test (*) p < 0.0001.

The presence of intact prophages and defence systems throughout the genomes correlated with a -0.39 dPR to increased phage resistance (Figure 6, Table S1). The tools used to detect defence systems offered different outputs, with some systems like Uzume or RloC being only detectable with one of the tools, and different classifications of the same CDS inside the same family (Abi2 or AbiD). Regarding CRISPR systems, a total of 167 different spacers were detected, out of which 20 showed hits against the viral genomes with varying degrees of specificity. Most hits affected all phages and none affected one phage specifically. In addition, in none of the viral genomes did we find any identities with anti-CRISPR (Acr) proteins.

**Figure 6.**
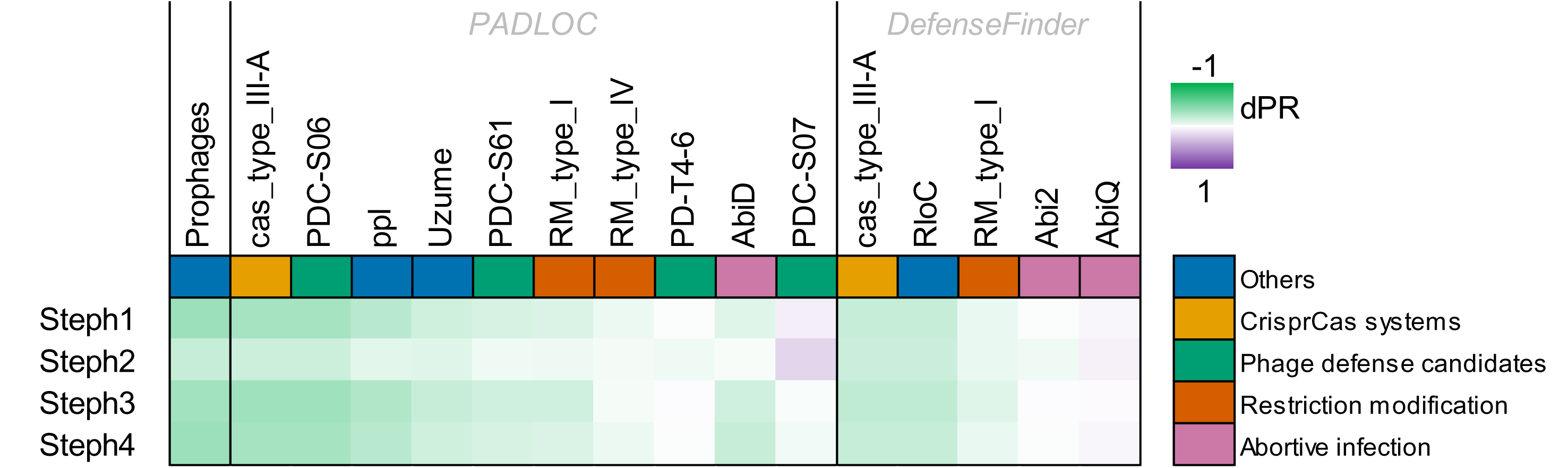
Potential association of bacterial defense systems with resistance to each of the phages. Only genes present, and absent, in at least 8 bacterial isolates are included. Color intensity determined by differential probability of resistance (green - correlated with resistance, and purple – correlated with susceptibility). Defense systems evaluated using PADLOC and DefenseFinder are shown.

Finally, analysis of the genomes of susceptible/resistant bacteria through GWAS and the posterior Bonferroni correction showed that resistance to Steph1, Steph3 and Steph4 was linked to the presence of genes *icaD*, *icaB*, *icaC*, *icaR*, *icaA* and *sbcC*. This showcases the relevance of the *ica* operon, involved in biofilm formation, and the *sbcCD* complex, involved in defence against phages carrying long palindromes^72–74^. Steph phages carried 210-218 palindromic sequences of up to 10 bp, of which a varying percentage (30-81%) were found in at least other Steph phage. Despite the correlation between the presence of the *ica* operon and *sbcC* with phage resistance in these genes, no genes were associated with increased susceptibility or resistance to phage Steph2 (Table S1).

## Discussion

The host range of phages is a crucial characteristic for understanding phage-host interactions and implementing phage-based applications to control bacterial infections. Phages with a broad host range can infect a larger number of bacterial isolates, increasing the likelihood of targeting the specific bacteria causing an infection. In this study, we isolated four closely-related phages infecting a large collection of clinical and veterinary *S. epidermidis* isolates, which highlights their potential as an alternative against clinical or veterinary pathogens. These broad-range phages belong to the *Herelleviridae* family and the *Sepunavirus* genus^75,76^. *Herelleviridae* phages, both from the *Sepunavirus* genus and others, have been used in phage therapy before with overall clinical improvement^33,77,78^. Since the use of antibiotics in agriculture and veterinary settings is highly regulated in Europe given its contribution to antibiotic resistance^79^, phage therapy might be a promising therapeutic tool in this setting.

Genomic analysis revealed the absence of integrases in all phages, supporting a strictly lytic lifestyle. However, at least one transposase was detected for Steph3 and Steph4 according to structural based predictions of coding sequences, and all isolated phages encoded one hemolysin. Phages containing transposases are not recommended for phage therapy given their potential to mobilize host genes (e.g. virulence factors and AMR related genes) and unforeseen effects that may result from transposition of DNA (for example, integration into regulatory elements)^80^. However, it is possible to inactivate or delete them before using these phages in a therapeutic setting. In addition, hemolysins are often found in phage genomes, as an identical sequence was found in the genome of phage BE06, previously used to treat musculoskeletal infections^78^.

Previous work highlighted the necessity of identifying host range determinants of staphylococcal phages^81^. In this work, GWAS analysis showed a correlation between the presence of genes *icaR*, *icaC*, *icaD*, *icaB*, *pgaC* (*icaA*) and *sbcC* and a protective effect against phages Steph1, Steph3 and Steph4. Biofilms protect bacteria against external threats, such as phages and antibiotics, by creating a physical barrier and regulating their metabolism^8,14,73^.

Polysaccharide intercellular adhesin (PIA) serves as the main structural component of *Staphylococcus epidermidis* biofilms, though PIA-independent biofilm formation has also been reported^82^. This exopolysaccharide mediates critical functions in biofilm development, promoting cell aggregation and providing protection against host immune responses^73^. The production and modification of PIA depend on the *icaADBC* operon, in which *icaA* encodes the poly-β-1,6-N-acetyl-D-glucosamine synthase responsible for PIA polymerization, while *icaD* enhances IcaA’s enzymatic activity^74,83^. The *icaB* gene product modifies PIA through de-N-acetylation, and *icaC* facilitates extracellular export and polymer elongation for proper matrix integration. Expression of this operon is tightly regulated by *icaR*, ensuring PIA production occurs under appropriate environmental conditions^74,83^. Interestingly, previous studies have demonstrated a synergistic effect between depolymerases and phages, which is hypothesized to result from the dispersive action of depolymerases on the biofilm matrix^84,85^. The addition of depolymerases capable of degrading PIA could be of particular interest where *icaADBC* has been correlated with phage resistance^73,83,84^. The predicted *icaADBC*-mediated resistance could be influenced by the use of a permissive, non-biofilm former isolate for phage hunting, which would be adding a bias towards phages unable to infect biofilm formers. However, broad-range phages were able to infect strong reference biofilm formers like *S. epidermidis* ATCC 35984 (RP62A). The presence of the *sbcC* subunit was also found to correlate with phage resistance. This nuclease forms part of the *sbcCD* complex, which is implicated in chromosome replication and interfering with infection of phages carrying long palindromes in their genome^72,86,87^. Regarding susceptibility to Steph2, no significant hits were obtained through the GWAS, perhaps due to the low number of infected strains. Another option would be that the main determinant of susceptibility against Steph2 is not determined by an annotated gene product, and instead by a sequence-dependent bacterial defence system, or other unknown mechanisms.

Prophages can confer resistance to other phages while integrated in a bacterial genome mainly through superinfection exclusion. Thus, a lysogenic bacterium can be more likely to survive in an environment with lytic phages, allowing the prophage to continue exploiting the bacterium to their mutual benefit^88^. Multiple systems have been described to this end, including preventing genome injection, using CRISPR systems or inducing abortive infection (ABI)^88–91^. Here, we showed that presence of at least one intact prophage was found to correlate with a lower chance of infectivity for the phages. Aside from prophages, the presence of different genes belonging to the ABI, RM and CRISPR-Cas systems in the bacterial genomes was also found to correlate with increased phage resistance. Had the genome of Steph2 contained CRISPR sequences not present in the other phages, this could have been responsible for the reduced host range of Steph2; however, no such CRISPR sequences were found. The general correlation between presence of CRISPR systems and phage resistance could be explained by an indirect hit, but CRISPR does not tend to co-localize with other defence systems, as opposed to RM or toxin-antitoxin systems^88,92,93^. It is important to highlight that all the associations were based on correlations between presence and absence of genes, and further experimental research is necessary to determine whether the presence of these defence systems does affect phage susceptibility.

Upon examining the genomes, we also found differential regions among the phages. The most pronounced divergence of Steph2 against its relatives was located in a region around 42 kbp, which includes CDSs encoding MTases in all four phages. All MTases were orphan, meaning that they were not associated with a restriction enzyme. This hints at a possible role in evasion of bacterial RM systems, something previously described in other Gram positive species, allowing the phage to adopt a methylation pattern in order to carry out successful infection^35,94^. However, our data cannot allow us to correlate with the effect on the host range of the phages. Future experiments involving deletion of the additional domain or exchanging of the MTases may provide evidence for its contribution to escaping host resistance mechanisms.

Understanding the interplay between phages and their hosts is essential for comprehending the potential of phages as therapeutics. In this study, we report the isolation and characterization of four broad-range phages capable of infecting a clinical and veterinary panel of *S. epidermidis* isolates. The *ica* operon and the *sbcC* subunit correlated with resistance against our phages, emphasizing the role of biofilm formation and anti-phage systems in determining phage host tropism. Future research should investigate the relationship between the genes identified in this study and their role in determining the host range of these phages, as well as bacterial defence mechanisms against the phages.

## Acknowledgements

We thank Dr Geoffrey Foster, SRUC Veterinary Services, Inverness, UK for providing the veterinary isolates of *S. epidermidis;* Dr Maria Miragaia, ITQB, Universidade Nova de Lisboa for isolate Gre26; Dr Micael Widerström, Umeå University, Sweden for isolate 99-1352, and Professor Lindsay Hall, University of Birmingham, UK for providing additional *S. epidermidis* isolates. NUH, Nottingham isolates were obtained from the Nottingham University Hospitals Trust Pathogen Bank, Nottingham, UK. We also thank Raquel Martínez-Recio as a student phage hunter.

## Supplementary table legends

Table S1. Genomic context of the bacterial isolates used in this study

Table S1.1. Isolation and genome information available of the Staphylococcus epidermidis isolates used in this study.

Table S1.2. Sequencing information available of the Staphylococcus epidermidis isolates used in this study.

Table S1.3. Defense systems hits as detected by DefenseFinder and PADLOC, respectively.

Table S1.4. Diferential probability of resistance (dPR) values of defense systems against phages Steph1-4.

Table S1.5. Genome-wide association study of bacterial genes and phage resistance.

Table S1.6. List of palindromes detected in the genomes of phages Steph1-4.

Table S2. Genomic context of the phages isolated in this study

Table S2.1. CDSs of phages Steph1-4 with a predicted funcion associated to structural, tail, or depolymerase activity.

Table S2.2. Mutations detected and shared by Steph1, Steph3 and Steph4, while absent in Steph2.

Table S2.3. Mutations detected in Steph1, Steph3 or Steph4, but absent in Steph2.

